# Magellon – an extensible platform for cryo-EM data visualization, management, and processing

**DOI:** 10.1101/2025.06.09.658726

**Authors:** Behdad Khoshbin, Puneeth Damodar, Rupali R. Garje, Frank H. Schotanus, Nandagopal Nair, Lai Wei, Jan-Hannes Schäfer, Gabriel C. Lander, Michael A. Cianfrocco, Scott M. Stagg

## Abstract

Single particle cryo-electron microscopy (cryo-EM) has revolutionized structural biology by enabling high-resolution determination of macromolecular structures. However, the field faces challenges in data management, processing workflow integration, and software extensibility. We present Magellon, an innovative cryo-EM software platform that addresses these challenges through a modern microservices architecture. Magellon consists of an extensible backend with a web-based front end that we call Magellon Viewer. Together, these combine high-performance computing capabilities with an intuitive user interface, enabling researchers to efficiently process and analyze cryo-EM data. The platform’s distinguishing features include a plugin-based architecture, distributed processing capabilities, comprehensive monitoring systems, and a novel approach to data organization and visualization. A key philosophy of the approach is that the Magellon backend provides a platform that uses robust industry-standard libraries to orchestrate computational tasks while offering users and developers flexibility in selecting the computational resources for performing calculations. Magellon represents a significant advancement in cryo-EM software infrastructure, offering flexibility, scalability, and extensibility while maintaining ease of use.

## Introduction

### Cryo-EM and automation

Automation has played a pivotal role in the explosive growth of single particle cryogenic electron microscopy (cryo-EM) and the accessibility of the technique to investigators who are not experts in electron microscopy. Automated tools and software can be found in all stages of cryo-EM structure determination, including specimen preparation^1,2^, data collection^3,4^, and image analysis^5–9^. The development and implementation of such tools has had a transformative impact on the field of structural biology, as they render the complex process of single particle cryo-EM structure determination more tractable, repeatable, and systematic.

The increased use of cryo-EM structure determination has been accompanied by an expanded collection of academic software for data processing, which continues to grow^10–12^. This growth has been accelerated by the rise of machine learning in computer vision, which has been integrated into nearly all aspects of the cryo-EM image analysis pipeline^13–16^. The software packages developed to address the various steps of data processing can be classified into three primary types: (1) comprehensive, all-in-one packages such as cryoSPARC^7^ or EMAN2^17^, where each processing step has been coded individually in the software package environment; (2) hybrid packages, such as Scipion, Relion, Appion or *cis*TEM^9^, which integrate both native algorithms and external software through wrapper interfaces; and (3) customized, bespoke data processing pipelines where individual investigators install the stand-alone programs and write convenience scripts to shuttle data between them. The advantage of the all-in-one packages is that they are relatively easy to install and learn to use, but have limited flexibility and extensibility. In contrast, bespoke pipelines, while highly customizable, can be complex to set up and difficult to manage. Hybrid software packages aim to combine the advantages of both approaches by providing a standardized interface that integrates both native algorithms and external software.

Hybrid systems and bespoke pipelines face several technical architecture challenges that affect workflow efficiency and the longevity of the software. One major problem is the complexity of software integration, as these systems often struggle to seamlessly integrate multiple processing tools and algorithms. Within the same vein are issues with data organization - when databases are incorporated into a software package, there is usually a lack of robust methods for portable data organization, which creates problems for inter-facility transfers. Scalability constitutes an additional limitation, with systems often unable to smoothly scale from personal workstations to large computing clusters. Finally, introducing new functionality can present considerable barriers, as integrating new software or algorithms typically demands comprehensive knowledge of the underlying codebase.

We present here a new hybrid software package, Magellon, an extensible platform designed to enable intuitive, point-and-click data viewing, organization, and analysis. Notably, the platform is open source, flexible, and designed for cross-compatibility across different operating systems. Magellon employs a microservices architecture to enable independent scaling of components, while its plugin-based design facilitates seamless integration of new algorithms without modifying core code. Considerable effort was devoted to the design of Magellon’s backend, making it deployable and consistent across different computing environments, including Windows, Linux, and MacOS. Additionally, Magellon employs industry-standard libraries and modern protocols to enable installation on personal laptops while maintaining scalability to data center-sized infrastructure. As a demonstration of Magellon’s capabilities, we developed the Magellon Viewer - a web-based, interactive frontend built with React JavaScript that leverages the Magellon backend. The platform’s modular design enables research teams to customize or replace components, such as the Viewer, with specialized interfaces tailored to specific needs, or even develop alternative frontends for particular applications. With potential for future expansion through micro-frontends, Magellon is positioned to become a critical component of cryo-EM cyberinfrastructure, supporting both individual researchers and large research groups.

## Design

### Design principles

The foundational philosophy behind Magellon is that it should be a platform-as-a-service (PaaS) tool for cryo-EM data processing and algorithm deployment. The architectural core of Magellon is structured to handle database organization, job submission, and user accounts. Additionally, Magellon offers extensible tool implementation through a well-defined plugin/toolbox interface in addition to enabling portability for data import/export. The combination of these tools establishes a foundational platform that can be used for data viewing and processing in a web-tool named the “Magellon Viewer”. This design ensures that training and maintenance remain manageable while promoting extensibility, thereby enabling the broader scientific community to contribute features and enhancements so that development is not limited to the core Magellon developers.

### Magellon Backend

The Magellon backend comprises a central data orchestration framework and a set of plugins, functioning similarly to a microservices environment where each component operates independently (**Fig. 1**). This design offers flexibility, scalability, and maintainability. The backend manages primary tasks, schedules jobs, and handles communication between plugins, while the plugins encapsulate specific algorithms that function as independent services. Magellon was designed to use resources available either on the host computer or across a network by leveraging representational state transfer (REST) programming. Using the REST framework, tasks are allocated to different workers that can reside on any computer accessible over the network with the Magellon software installed. To further the robustness of our backend infrastructure and accommodate ease-of-use for developers aiming to extend or interact with Magellon tasks, we developed a Magellon application programming interface (API). The REST API framework was written using FastAPI (https://fastapi.tiangolo.com) and Swagger (https://swagger.io) (**Fig. 1**), and enables task requests to be transmitted over a network. FastAPI provides for separation of controllers for each purpose (e.g. separate controllers web, database, processing, etc.) and enhances code organization and readability, while Swagger provides an interface for API documentation and testing.

**Figure 1.**
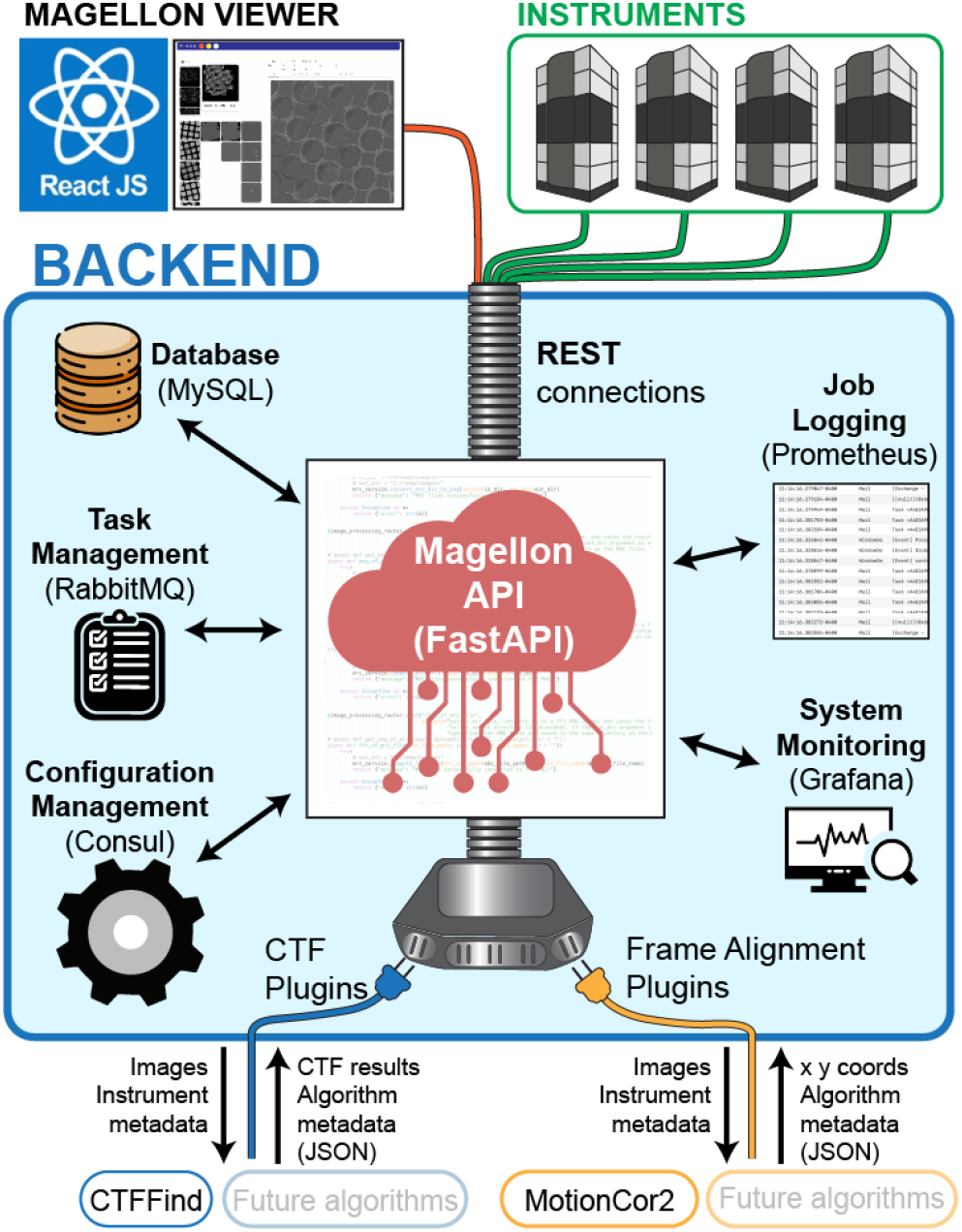
Magellon REST computational architecture and API Framework. The heart of Magellon is the Magellon API which utilizes industry standard libraries and frameworks that altogether form the Magellon Backend. The Magellon Viewer Frontend and plugins connect to the backend through standardized outlets that enable extensibility of the platform.

To effectively manage the diverse processing tasks that could be carried out within the Magellon interface, which could range from seconds to hours of compute time, a robust task manager was implemented using RabbitMQ (https://www.rabbitmq.com) for message queuing and management (**Fig. 2**). RabbitMQ is an open-source message broker with established reliability and efficiency in handling asynchronous communication between distributed systems. This integration ensures that tasks are efficiently scheduled, queued, and executed across various computational resources, optimizing Magellon’s performance and scalability.

**Figure 2.**
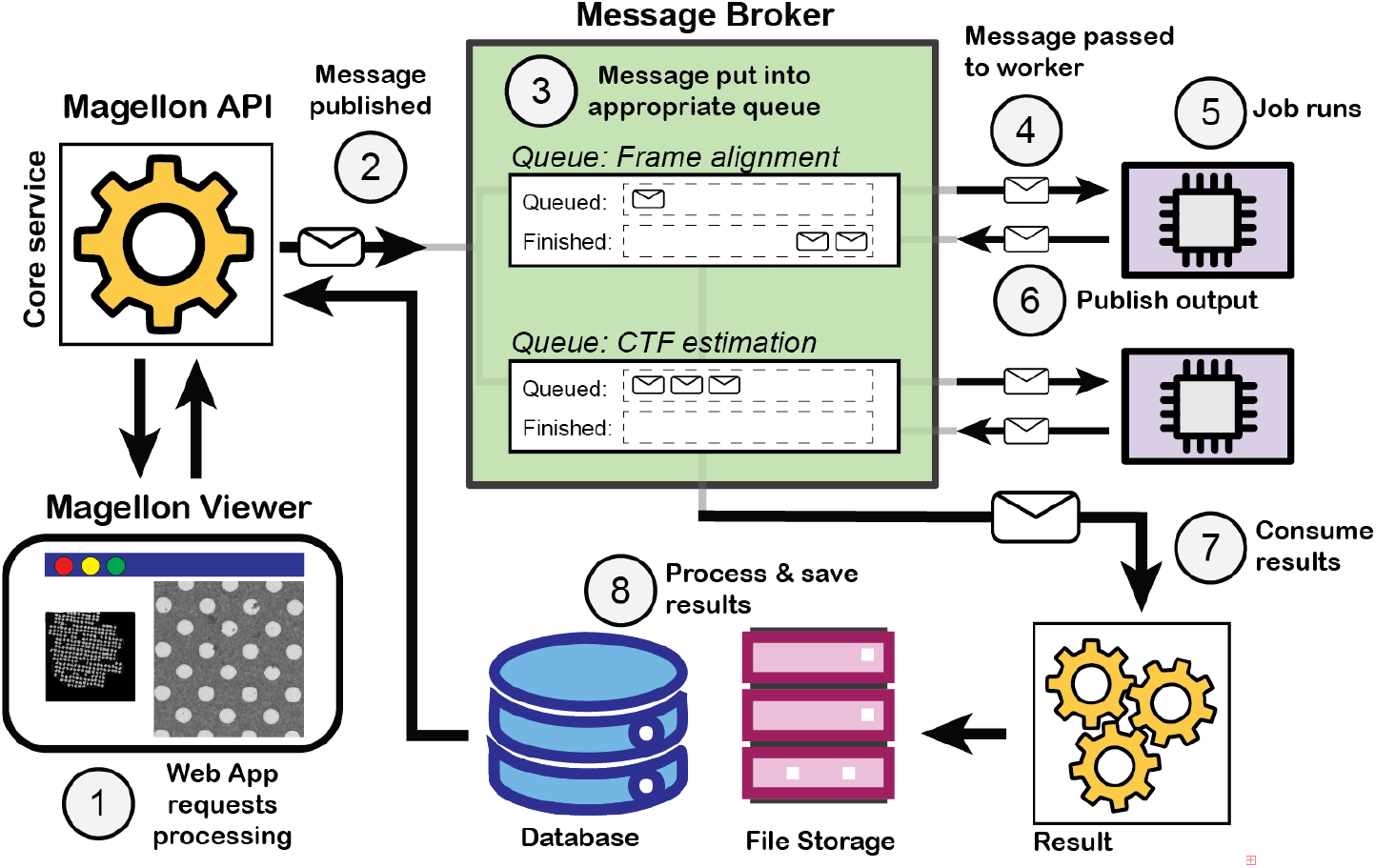
Task management in the Magellon backend. 1) Tasks are initiated from the Frontend and passed to the Magellon API, 2) Messages are published to the RabbitMQ message broker. 3) Messages (i.e. job requests) are placed in a job queue. 4) Jobs are passed to a data processing worker, where 5) the job runs. 6) When the job completes the outputs are published back to RabbitMQ. 7) RabbitMQ routes the results to the results processor that 8) inserts metadata into the Magellon database and saves the results to the appropriate paths in the file system and notifies the Magellon API of completion.

Maintaining consistent and validated data exchange between different REST clients is crucial to Magellon’s efficiency and the integrity of analyses. To this end, Magellon employs Pydantic models (https://docs.pydantic.dev) for data structuring and validation. Pydantic is a Python library designed for data parsing and validation that integrates seamlessly with FastAPI and Swagger frameworks. Pydantic ensures that incoming data adheres to the defined schemas and formatting requirements, thereby minimizing the risk of errors occurring during processing interactions. For example, in Magellon there is a Pydantic model for images (ImageDtoBase) that defines all of the metadata and types (e.g. stage position, image shift, spot size, beam intensity, pixel size, etc.) that are expected for a TEM image that is being imported. Thus, data import is standardized regardless of what instrument or software used for data acquisition. By standardizing data formats in this manner, Magellon facilitates streamlined interactions within the platform, simplifying and standardizing the development process.

### Frontend: Magellon Viewer

The Magellon Viewer is entirely web-based to maximize accessibility, ease of interaction, and exposure of users to their data and metadata while simultaneously leveraging the suite of backend Magellon tools. The web navigation is designed to be intuitive, being primarily image-based with a hierarchical organization of the low- and high-magnification images that comprise a dataset. Relevant metadata are prominently displayed and accessible, ensuring that users can access any associated information with a dataset (e.g. microscope and detector settings). Web pages are generated dynamically, and interactivity is achieved through JavaScript using the React framework (https://react.dev), which ensures a responsive and seamless user experience.

Interaction between the frontend and backend is facilitated through RESTful API calls. When a user interacts with the web interface, such as by selecting a dataset or processing task, the frontend sends requests to the backend via the Magellon API. Processed data are communicated to the frontend, updating the interface in real-time to enable navigation of large datasets, track the status of processing jobs, and review results instantly. Associated metadata are displayed in a structured and user-friendly format, often alongside corresponding images. For instance, metadata such as acquisition parameters (e.g. keV, magnification) and processing results (e.g. particle selection, contrast transfer function (CTF) estimation) are integrated into the display. The metadata are dynamically loaded from the backend, ensuring that any updates or changes made during data processing are immediately reflected in the interface, which is particularly important for users to make informed, real-time decisions based on the incoming data.

### Magellon database structure

To handle the complexity of metadata associated with a cryo-EM data collection experiment, Magellon employs the relational database MySQL (https://www.mysql.com) for efficient storage and retrieval. Many concepts were taken from the database architecture developed previously for the data acquisition software Leginon^3^, but the Magellon database is greatly simplified to facilitate easier data export and transfer

### Magellon adaptors to enable cross-software compatibility

Magellon was developed to be agnostic to the software used for data collection. Data and metadata are ingested and assigned to unified metadata fields so that the import process is the same for all data collection packages. This approach is accomplished with a Magellon translator layer that populates defined metadata fields and inserts them into the Magellon database. Pieces of code, called “Adapters,” read metadata from different data collection packages and interface with the translator to enable imports from various software sources. Adapters have been written for the data collection software packages Leginon^3^ and Thermo Fisher Scientific EPU, and adapters for SerialEM^4^ and Gatan Latitude are currently under development (**Fig. 3**). Additionally, Magellon import/export utilities have been written to enable data transfer between different sites.

**Figure 3.**
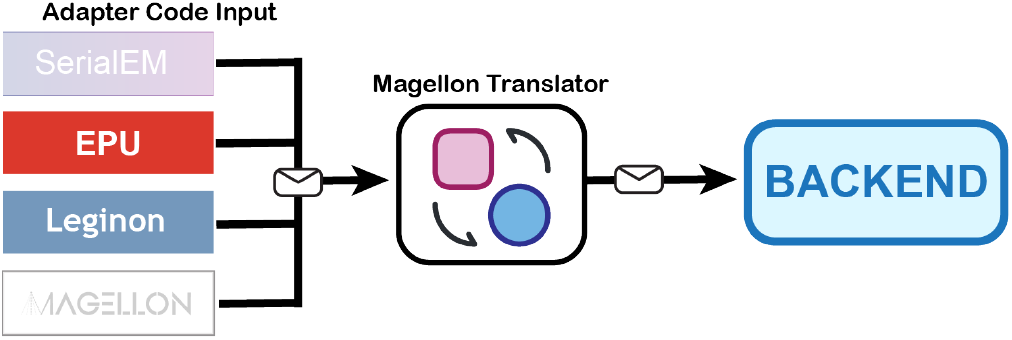
The Magellon translator. Adapter code unifies metadata from multiple different input sources and inserts the data in a common format into the Magellon database. between data collection sites. A potential problem in designing a database where the metadata can be transferred between different physical locations is that a mechanism is needed to ensure that primary keys are not duplicated. Thus, to avoid primary key conflicts, we use universally unique identifiers (UUIDs) as primary keys. This guarantees that all systems have globally unique IDs for their image records, allowing different databases to be merged readily into one database without concerns about conflicts. There is no built-in UUID column type in MySQL, thus UUIDs are stored as binary(16) by removing four characters and storing each two hexadecimal numbers in a single byte. The full database schema is available for viewing on the Magellon website (http://magellon.org).

### Magellon’s plugin interface offers flexible incorporation of software tools

Magellon is designed with an open architecture that promotes extensibility, facilitated by a robust plugin system that streamlines the testing and integration of new algorithms (e.g., CTF, Fast Fourier transforms (FFTs), particle picking, frame alignment, etc.). Built on a microservices architecture, each data processing algorithm exists as a distinct plugin or microservice, offering significant advantages over monolithic designs. Plugins can be independently developed, tested, published, and shared, while also being deployable on separate systems or via containers with isolated dependencies. Each plugin functions as a self-contained FastAPI project encapsulating a single algorithm, ensuring modularity and independent development cycles. This approach not only enhances scalability but also simplifies deployment through containerization technologies like Docker and orchestration tools like Kubernetes (https://kubernetes.io).

Magellon’s plugin architecture draws inspiration from the Akka framework’s (https://akka.io) actor model, where each plugin operates as an independent actor within the system. Plugins communicate with the backend exclusively through message-passing via RabbitMQ, maintaining dedicated message queues that enable asynchronous task processing, state isolation, and operation encapsulation. This approach offers significant advantages: it prevents individual plugin failures from affecting the entire system; hierarchical supervision automatically restarts failed plugins; and state preservation mechanisms safeguard data during interruptions. The design ensures location transparency, allowing plugins to be relocated or scaled without system disruption. By utilizing message-passing for parallelization, the architecture efficiently manages resources and minimizes bottlenecks while enabling horizontal scaling and dynamic resource allocation. Further, each plugin’s isolated memory space ensures optimal memory management and reduces conflicts. These architectural choices make Magellon’s plugin system inherently extensible, scalable, and resilient, enabling cryo-EM algorithm developers in the community to deploy their specialized algorithms as Magellon plugins without concerning themselves with peripheral challenges like data management, system control, pipeline configuration, or parallelization. Developers can thus focus exclusively on algorithm development while Magellon handles the infrastructure.

### Flexible Magellon deployment for individual users or large facilities

Magellon was designed with a “deploy anywhere” philosophy, emphasizing ease of installation while maintaining flexibility for diverse computational environments. This approach directly addresses the software compatibility and installation challenges commonly faced in hybrid computing infrastructure projects. The deployment architecture supports both simple single-machine installations for individual researchers and complex distributed setups for large facilities, achieved through two complementary approaches: Docker Compose for containerization and a custom Command-Line Interface (CLI) for management. Magellon’s deployment architecture accommodates three primary scenarios: single-machine, distributed, and hybrid deployment. In the single-machine deployment regime, all components of Magellon run on one computer while maintaining isolation through containerization, requiring minimal configuration yet providing full platform functionality. For distributed deployment, Magellon components can be installed across multiple networked computers, with each component placed strategically to utilize specialized hardware (such as GPU clusters) while maintaining seamless system-wide communication. In hybrid deployment installations, Magellon can bridge local resources with cloud infrastructure, allowing researchers to maintain core functions locally while dynamically expanding processing capabilities to external resources as needed. This flexibility ensures that Magellon can adapt to computational needs in a cost-effective manner, enabling dynamic scaling, burst processing for large datasets, and geographic distribution of workloads. Together, these flexible deployment options ensure that Magellon can adapt to a wide range of user needs and institutional scales.

Magellon’s modular architecture enables distributed deployment, where each component can be installed on separate machines across a network. This distribution capability allows components to be strategically placed on hardware optimized for their specific requirements, such as GPU-accelerated nodes for computation-intensive plugins or high-memory servers for database operations. The system’s components automatically discover and communicate with each other through standardized APIs and message queuing, functioning cohesively as an integrated platform, regardless of physical location. This functionality is facilitated through the use of Consul, which is an open-source networking and service discovery tool for dynamic, distributed environments. Consul provides a registry where services can register themselves and discover other services, streamlining communication in microservices architectures.

This architecture provides both forward compatibility and scalability. Research facilities can manage their project uniformly through a single Magellon instance distributed across their infrastructure. The platform supports both vertical scaling (adding more CPUs, GPUs, or storage to existing nodes) and horizontal scaling (incorporating additional computers or compute nodes into the system) without requiring architectural redesign. Moreover, with proper network configuration, Magellon enables geographic distribution of resources. Conceivably, a facility could maintain data storage in one location (e.g., Florida) while leveraging idle GPU resources at another institution or the cloud, creating a federated computing environment that maximizes resource utilization across sites.

There are two mechanisms for deployment, depending on the computational needs and expertise of the end user. For single workstation deployment, individual components have been containerized using Docker. A single Docker “compose” command can install all of the necessary dependencies onto a workstation of interest. For developers or on distributed systems, there is a command-line interface (CLI) tool that was developed as a lightweight standalone executable. The CLI handles deployment, setup, and management of both the Magellon core system and its plugins. Its design is platform-agnostic and has been tested on Windows, Linux, and macOS. A single command “magellon install app” initiates the entire installation process. This command handles all the necessary environment setup and configuration, ensuring that users can get started with minimal effort and eliminating the need for manual configuration steps. The CLI simplifies the development process with the commands “magellon development setup” and “magellon development load,” which automatically configure the environment and load necessary dependencies. Together, these deployment tools provide a highly accessible and flexible entry point for users and developers, lowering barriers to adoption and promoting community-driven expansion of Magellon’s capabilities.

Plugin management is also handled with the Magellon CLI. Developers can create and deploy plugins using the commands “magellon plugins create init,” “magellon plugins create myctf,” and “magellon plugins deploy myctf.” Likewise for end users, plugin discovery and installation are handled with the command “magellon plugins list -a -s:author,” which allows users to browse available plugins, while “magellon plugins install myctf@latest” enables seamless installation of specific plugins. The CLI makes it straightforward for users to expand Magellon’s functionality with minimal effort, without needing deep technical knowledge.

## Materials and Methods

### Data collection for demo

A small cryo-EM dataset was collected to demonstrate the functionality of Magellon for data import, management, and processing. Mouse heavy chain apoferritin from a plasmid gifted by Dr. Masahide Kikkawa^18^ was expressed and purified as described previously^19^. UltraAuFoil R1.2/1.3 300 Mesh grids were plasma cleaned for 50 seconds at 7 Watts with 25% oxygen, 75% argon (Gatan Solarus 950). Cleaned grids were loaded onto Vitrobot Mark IV at 100% humidity (4 ^*°*^C). 4 µL apoferritin was applied to the grid which were blotted for 1 second at 0 force before being vitrified in liquid ethane. Data were collected on a Titan Krios microscope (ThermoFisher Scientific) operated at 300 kV with a 50 µm C2 aperture and 100 µm objective aperture and equipped with a DE Apollo direct detector. Images were acquired using Leginon^3^. For high-magnification exposures, the Apollo was set in 8k × 8k super-resolution mode at a magnification of 59,000× with a super-resolution pixel size of 0.395 Å. The movie frame rate was fixed at 60 frames per second.

## Results

### Deploying Magellon and Importing Data

As described above, Magellon can be installed and run using Docker or installed manually by setting up the required dependencies. Detailed instructions for both methods are available on the Magellon website (http://magellon.org) or the ReadMe file included with the source code. To help new users become familiar with Magellon, the code is accompanied by a test dataset that can be used to explore Magellon functionality. This small test dataset of vitrified apoferritin has been deposited in the Electron Microscopy Public Image Archive (EMPIAR) under the accession number EMPIAR-11254. The entry includes all 310 images associated with the dataset, including low-magnification images and 218 high-magnification exposures with accompanying movies.

To streamline data handling operations, we utilized Adaptors within Magellon to build a web tool for Magellon import/export, which enables users to import the test dataset seamlessly into the Magellon Viewer. We tested import on a variety of computers and platforms (**Table 1**). The test data includes both integrated micrographs and movies. During the import process, Magellon ingests associated metadata, creates image files and thumbnails for every image in PNG format, pre-calculates power spectra for every image, and estimates the CTF for every image with a pixel size smaller than 5 Å /pix. Since the distribution is containerized using Docker, the processing operations are platform-independent and the Linux binaries that Magellon wraps can be executed seamlessly on a variety of Linux distributions, Mac OS, and Windows: essentially any OS for which Docker is compatible. The program “MotionCor2” serves as the default Magellon frame alignment plugin, so the frame alignment jobs will only execute on computers that are equipped with a compatible Nvidia GPU. However, the Magellon import process will proceed regardless of the presence of a GPU, enabled by the robust fault tolerance of RabbitMQ, which allows for the execution of jobs independently without the failure of one job propagating throughout the system.

**Table 1.** Magellon data import statistics.

| Platform | Chip | GPU | RAM | Processing time | Frame alignment |
| --- | --- | --- | --- | --- | --- |
| Mac OS iMac | M1 | None | 16 G | 19 min | None |
| Linux | Xeon W | GeForce RTX | 256 G | 1h 45 min | Yes |
| Linux | Xeon Silver | GeForce GTX | 64 G | 5h 59 min | Yes |

We recommend that new users of Magellon become acquainted with the software by importing the test dataset and following the step-by-step demonstration included in our user guide. This demo provides an overview of Magellon’s key features and helps users understand how to integrate their own data into the system.

### Interacting with Magellon Viewer

The first step for starting a session with Magellon Viewer is filling out a web-based metadata form containing information about the user’s data. Upon completion, the user submits the import job, which triggers a series of customizable processes including metadata import, transfer of raw data to its final location, frame alignment, and CTF estimation. Import can occur concurrently with data collection, and the data processing occurs on-the-fly.

The Magellon Viewer Graphical User Interface (GUI) was designed to maximize the visibility of the data and metadata. The interface is organized into four main sections: an atlas viewing section at the top left, a hierarchical image browser below the atlas section, a metadata display section on the upper right, and a large interactive image viewer below the metadata section (**Fig. 4**). Users are presented with images representing the sample at every magnification used during data collection. Clicking on an atlas image reveals all the associated low-magnification images in the hierarchical browser, providing an overview of data acquisition at different scales. Colored bounding boxes highlight images that served as the basis for targeting higher-magnification acquisitions, visually guiding users through the imaging hierarchy.

**Figure 4.**
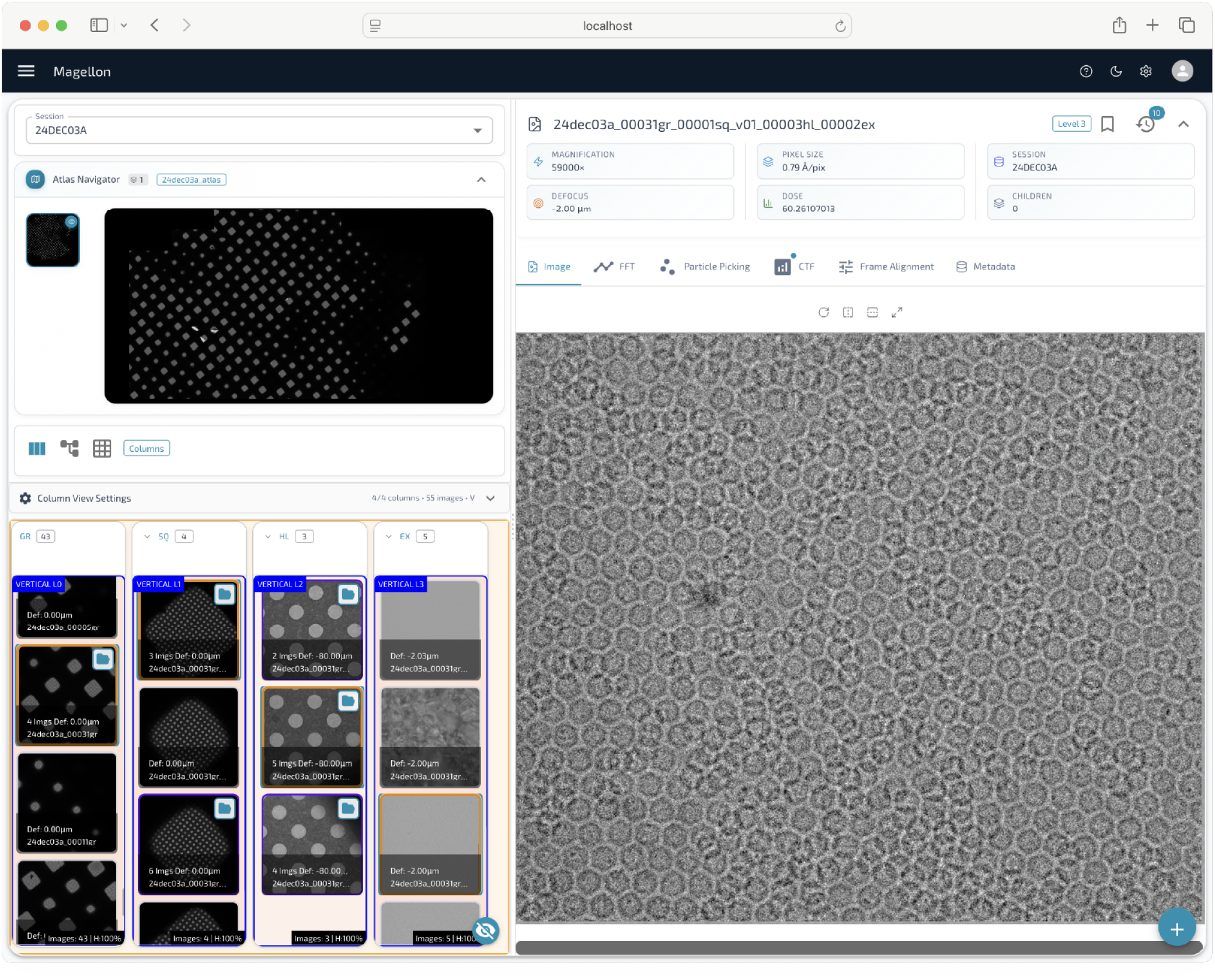
The Magellon viewer. The viewer shows atlases for a given session at the top left. A hierarchical image navigator is featured below which allows users to navigate between lower magnification images and their higher magnification children. The center features an interactive viewer with associated metadata.

### Plugin development: CTFFIND4 and MotionCor2

Two plugins have been developed for Magellon that serve as examples for future plugin developers. These two plugins carry out the initial image analysis processes that are universally performed on all cryo-EM data – frame alignment/dose-weighting and CTF estimation. Each type of image analysis process in Magellon is designated to a specific plugin category with defined metadata that are provided from Magellon to the plugin. Likewise, outputs are generated with the expected syntax, format, and in an organized fashion. The inputs and outputs are managed by specific queues in the Magellon process manager controlled by RabbitMQ. Since inputs, outputs, and metadata to and from the plugin are in a predefined format, the Magellon backend can handle queries, insert metadata into the Magellon database, and generate organized output files. As Magellon continues to develop, more types will be developed (e.g., particle picking, image assessment, etc.). This approach grants flexibility to developers, so that standardized inputs and outputs can be directed to any new algorithm, eliminating the need to rebuild or reconfigure the entire system.

When a user selects an image from the hierarchical browser, a full-resolution version is displayed in the interactive image viewer. This viewer offers a range of built-in tools, including the ability to display precomputed FFTs, a measuring tool for estimating distances or dimensions, and an interactive particle picker for manual or assisted particle selection. Key metadata, such as contrast transfer function (CTF) estimation results, magnification, and pixel size, are prominently displayed above the viewer for immediate reference. Full metadata records, including all recorded microscope parameters, camera settings, and processing metadata, are readily accessible through an expandable tab, enabling users to intuitively retrieve detailed information about each image and the corresponding acquisition conditions. This integrated, data-centric design ensures that users can efficiently navigate complex datasets, assess image quality, and manage data collection strategies within a single unified platform.

The CTF estimation plugin was developed using the widely adopted CTFFIND4 program^20^. To manage task submission and execution, we established two RabbitMQ queues: an input queue for receiving processing requests and an output queue for returning results. The task is generated based on user-defined input parameters and submitted through the input queue. The input queue consumer retrieves each task and initiates the CTF estimation process, which comprises two main phases: CTF estimation and CTF evaluation. During the estimation phase, a command-line instruction for CTFFIND4 is dynamically generated using the input parameters and executed via a subprocess call to the CTFFIND4 binary. If an error occurs during execution, an error response is generated and returned; if successful, the outputs, including CTF parameters and diagnostic images, are captured for further evaluation. Following estimation, a secondary evaluation phase is conducted using quality metrics originally implemented in Appion^21^. The outputs from CTFFIND4 are subsequently analyzed, and all relevant metadata needed for CTF evaluation are collected and stored. Each request is assigned a unique directory, ensuring organized storage of output files and associated metadata. Outputs are formatted consistently across all plugins, then passed to the output queue for downstream processing. The result processor plugin subsequently retrieves and organizes the outputs, saving files to designated folders and recording metadata entries in the database. Outputs are classified into relevant categories—such as CTF or frame alignment—allowing seamless integration with the broader data management infrastructure. To ensure platform independence and facilitate deployment across diverse computing environments, both CTFFIND4 and the associated CTF evaluation code were containerized using Docker.

The frame alignment plugin follows a similar architecture to the CTF plugin and was developed using the MotionCor2 program^22^. Like the CTF pipeline, frame alignment uses two dedicated RabbitMQ queues - one for input and one for output - to manage task submission and result handling. Because MotionCor2 leverages GPU acceleration, special care was taken to ensure that all GPU drivers, libraries, and dependencies were correctly installed and compatible with the version requirements of the software. Upon receiving a task from the input queue, the plugin dynamically generates the appropriate MotionCor2 command using metadata retrieved from the Magellon database and executes the command via a subprocess. Log files, aligned micrographs, and associated outputs generated during processing are systematically collected, organized into structured folders, and registered within the database via the output queue. To promote platform independence and reproducibility, MotionCor2 and its supporting components were containerized in a dedicated Docker image, so that the plugin runs consistently on any machine equipped with an appropriate Nvidia GPU.

Integration of all processing plugins into the Magellon core service in this manner ensures seamless operation within the broader platform. Task creation for plugins is automated by querying the necessary input metadata from the Magellon database and dispatching processing tasks to the appropriate input queues. The core service coordinates the execution of plugins by managing task scheduling, monitoring process status, and ensuring that outputs are appropriately stored and categorized. This modular and standardized architecture not only streamlines cryo-EM data processing but also facilitates extensibility, enabling straightforward incorporation of new algorithms and tools into the Magellon framework.

### Magellon’s swagger interface: a web-based description of tools within Magellon

To enhance transparency, flexibility, and ease of customization, Magellon provides users with direct access to backend services through an integrated Swagger interface. Swagger is an open-source tool for visualizing and interacting with RESTful APIs in a user-friendly, web-based format. Rather than requiring users or developers to interact directly with the underlying codebase, Magellon exposes key backend functions through Swagger, which enables users to query data, launch processing jobs, manage configurations, and automate workflows directly through a point-and-click web interface.

Users who install Magellon through the demo Docker containers can access the Magellon Swagger interface by navigating to the website “http://localhost:8000/docs.” Through this portal, users and developers can view detailed documentation of available backend functions, including descriptions of required input parameters and expected outputs. For example, users can retrieve specific dataset metadata, submit processing jobs, or perform individual processing tasks. This design lowers the barrier for advanced usage and extension of Magellon, offering a straightforward pathway for users to explore backend capabilities and for developers to build custom features.

To introduce users to the Swagger interface, we implemented a standalone 2D lowpass filter function, which allows users to apply a lowpass filter to images already imported into Magellon or to new images uploaded by the user for on-the-fly filtering. Importantly, this example provides an accessible entry point for developers who want to begin building and testing their own image-processing algorithms within Magellon. Users can use this tool to selectively process individual images rather than the full dataset, showing how developers can quickly prototype and validate their routines in a lightweight, low-barrier environment. The lowpass filter plugin thus serves as a template for developers interested in contributing new functionality to the Magellon backend, demonstrating how custom processing functions can be exposed through the Magellon API and integrated into the larger workflow. Additional documentation and tutorials on how to use the Swagger interface, including examples for extending Magellon with new processing modules, are available on the Magellon website (http://magellon.org).

## Discussion

New algorithms are continually being developed for cryo-EM from many different sources, each with its own dependencies. Integrating these new algorithms into a given processing pipeline represents a significant challenge for both developers and end users. A key aspect of Magellon is extensibility, and we have developed a plugin infrastructure that enables investigators to develop and deploy new utilities for cryo-EM data processing. We have developed two initial plugins including one for CTF estimation and one for frame alignment that serve as examples for future development.

Magellon’s architecture aligns with a broader shift in how cryo-EM data collection and processing are evolving. While current tools such as Thermo Fisher’s “CryoFlow,” cryoSPARC Live, and modules built around Leginon and SerialEM data collection software enable users to preprocess data and adjust acquisition parameters in real time, Magellon takes this concept significantly further. We envision a seamless, decoupled workflow wherein data collection and analyses are no longer tied to the same physical location, but remain fully integrated operationally. Magellon’s microservice-based deployment model and data management capabilities uniquely position it to enable this vision. For example, a researcher could initiate a data collection session at a remote facility through the Magellon web interface. As micrographs are acquired, they would immediately appear in the researcher’s browser - without any visible indication that behind the scenes, raw data are being 1) temporarily cached at the facility, 2) automatically transferred to high-performance cloud storage, 3) registered in a globally distributed Magellon database, 4) processed by dynamically provisioned cloud compute resources, and 5) possibly being hosted at a geographically distant site, as the researcher visualizes the images through a web interface. The user remains shielded from the complexity of the underlying infrastructure, experiencing a unified and seamless workflow regardless of where the physical resources are located.

This location-transparent approach democratizes access to advanced cryo-EM capabilities, benefiting researchers in small laboratories or users of large research centers. Institutions without extensive local infrastructure could leverage cutting-edge microscopes and cloud-based computational power without costly investments in hardware that rapidly becomes obsolete. Collaborators at different institutions could simultaneously view and analyze datasets in real time, thanks to Magellon’s ability to ensure a consistent and integrated view of project status and data processing. For facility managers, the architecture offers flexibility and scalability: computational resources can be dynamically scaled to meet demand, with operational costs tied directly to usage rather than fixed infrastructure capacity. Moreover, processing jobs could be routed to regions with lower energy costs or available surplus compute capacity, improving both cost efficiency and environmental sustainability. Magellon’s current design, which separates data management, visualization, and processing components, lays the critical foundation for this future, where physical resource constraints are minimized and scientific inquiry can proceed with unprecedented fluidity and reach.

In the coming years, we anticipate that Magellon will evolve into a comprehensive cryo-EM data collection and viewing platform that will continue to integrate algorithms and software to enhance data acquisition efficiency. We plan to expand the plugin ecosystem, introducing additional tools for particle picking and image classification while simplifying capacity for user-based development through a dedicated CLI, reusable SDK, and robust documentation. A centralized plugin hub will facilitate sharing and collaboration among researchers, leading to new discoveries. The Magellon back-end sets the foundation for future data collection tools that will leverage its database-backed processing infrastructure, enabling integration with Magellon Viewer for real-time data visualization and assessment. Direct instrument integration will enable real-time control and data capture from the microscope and detectors, with the flexible infrastructure designed to accommodate both single particle as well as cryo-electron tomography data import, analysis, and visualization. The current job management system will mature into a fully automated cryoEM data collection engine supporting advanced features that can distinguish appropriate regions for data collection and associated event triggers. Users will also benefit from a responsive supervision dashboard accessible from both desktop and mobile devices, permitting remote monitoring of collection sessions. Finally, we aim to streamline cloud deployment with one-click AWS implementation, enabling efficient cloud-based data processing with intelligent resource management to optimize performance while minimizing costs.

## Acknowledgements

Funding for this research was provided by a National Institutes of Health grant R01 GM143805 to SMS, GCL, and MAC. The authors thank JC Ducom and Charles Bowman at Scripps for computational support, and members of the Cianfrocco, Lander, and Stagg lab members for beta testing and providing feedback on early Magellon releases.

## Author Contributions

The Magellon backend and frontend code was written by BK. CTF and Frame Alignment plugins were written by PD. The initial draft of the manuscript was written by SMS and BK, and subsequently edited by all authors.

## Data and software availability

The apoferritin dataset used for testing is available on the Electron Microscopy Public Image Archive under accession code EMPIAR-11254. Code is available on GitHub (https://github.com/sstagg/Magellon.git), and documentation is available on the Magellon Website (https://www.magellon.org/).

